# Uncovering the role that specific genes play in reachability properties of complex biological Gene Regulatory Networks (GRNs): The Arabidopsis thaliana flower development case

**DOI:** 10.1101/2023.05.30.542986

**Authors:** José Eduardo Chairez-Veloz, Elva Carolina Chávez-Hernández, José Dávila-Velderrain, Elena R. Álvarez-Buylla, Juan Carlos Martínez-García

## Abstract

The dynamic processes of multicellular organism development are regulated and coordinated by Gene Regulatory Networks (GRN’s). Therefore, a sustained effort to understand the dynamical properties of these modularly structured networks has shown the great utility of experimentally grounded and dynamically characterized discrete Boolean models, as an ideal formalism based on dynamical systems modeling tool for its qualitative description. Up to now, several low-dimensional Boolean GRNs have been proposed to recover gene activation configurations observed for specific cell types, and they have been validated via robustness and mutant analyses. Nevertheless, systematic studies that elucidate the role of individual genes implicated on transitions between given attractors in the context of morphogenetic patterns of development, are still very scarce. Such sort of studies is in fact quite relevant because genes belonging to a given GRN do not work in isolation. Indeed, they could interact with others GRN’s and/or with micro-environmental cues. Consequently, the structural specificities of the involved genes at the network level should be assessed in order to uncover the functional nature of the larger network involved. This is particularly meaningful when considering the role played by specific genes on transient dynamics related to cell fate specification. Following this idea, we propose here a computer-based analytical procedure intended to elucidate the role that specific genes play on the reachability properties of GRNs. As a structural property of a given dynamical network, reachability characterizes the attainability of specified attractors from given initial attractors as a consequence of the action of specific driving exogenous stimuli. Our proposal is based on algebraic systems approaches built around the Semi-Tensor Product (STP). We illustrate here our proposed procedure through the exploration of the reachability properties of the well-known Floral Organ Specification GRN of Arabidopsis thaliana (FOS-GRN), that recovers ten fixed-point attractors. Our findings suggest that there exist 79 inducible transitions among all possible pairs of attractors, with a suitable external Boolean control input over different well-characterized nodes of the network. Additionally, we found that such potentiality of these genes to produce attractor transitions is maintained by the continuous approximation model of the FOS-GRN, recovering not only qualitative but also useful quantitative information. Finally, we discussed the biological significance of our results and, even if we do not establish the specific molecular nature of the characterized exogenous control input, we concluded that *reachability analysis* can give us some important insights on the network level role that individual genes acquire by their collaboration with the GRN, becoming then targets in cell-fate decisions during development.

**Author summary:** Bringing to light the specific role that given genes play in gene regulatory networks is of particular importance, especially when it is necessary to quantify the influence of the environment on their dynamics. However, this becomes difficult by the fact that the genes in the network interact both nonlinearly and in the presence of feedback-based interactions. This requires then the development of methods of analysis that take into account such complex interdependencies. In this regard, Control Theory offers tools that allow characterizing the changes in the transition patterns between stable configurations (that is, cellular phenotypes) of the networks as a result of the presence of exogenous *stimuli*. In this work we propose a method based on the algebraic representation of small size gene regulatory networks, we first described in discrete Boolean terms, focused on delimiting the influence that specific nodes of the network play in the enhancement of transitions that define trajectories in the space of stable configurations of the state of expression of the genes involved. The proposed method uses the key control-oriented concept of *reachability* and is illustrated by the characterization of some induced morphogenetic trajectories that explain the development stages of *Arabidopsis thaliana* flower organs. Our proposal allows us to confirm that, in biological terms, the reachability analysis offers powerful tools to deepen the understanding of the interplay between the structure of specific gene regulatory networks involving both their own constitutive elements and including the networks with which they interact. This contributes to the understanding of biological development, which opens an access route to the exploration of the basic principles that associate the structure of gene regulatory processes not only with cell reprogramming and cell dedifferentiation, but also with dynamic processes underlying phenotypic plasticity and its evolutionary consequences. This is because the explanation of phenotypic change responses to environmental variability requires specifying how the constraints that govern developmental trajectories, potentially elucidated through reachability analysis, modulate the balance between phenotypic robustness and evolvability.

## Introduction

The continuing effort to understand how genes relate to the phenotypes of living organisms, leads to increasing volumes of experimental information resulting from the study of the complex biomolecular systems involved in cell differentiation as well as in morphogenetic dynamics. In this context, the use of mathematical modeling has become mandatory [1–3]. Gene Regulatory Networks (GRNs) models have been shown to be particularly useful to this end. Such complex biomolecular networks that underlie cell differentiation are mainly constituted by nonlinear interactions, presenting positive and negative functional feedback loops, which are critical for the establishment of their dynamic properties. In fact, recent studies have shown that the combination of such functional loops are crucial for the delimitations of the network’s stable configurations [4]. These stable configurations, referred to as network’s attractors, have been consistently interpreted as the cellular gene-expression profiles that emerge during differentiation. In general, the identification of developmental GRNs comes from the integration of well-curated functional data and the interactions of the genes involved. The characterization of these networks and their subsequent analysis has mainly focused on recovering observed gene activation configurations for studied cell types and their validation via robustness and mutant analyses, as well as new available experimental data. Such networks have, hence, recovered the qualitative and complex nature of regulatory modules involved in different aspects of plant and animal development [1, 5–8], and in some cases modules that are relevant for human development and biomedicine [9–11].

There have been a number of attempts to model developmental GRNs through formal approaches, i.e., linear models [12–14], systems of Ordinary Differential Equations [15] (ODEs), and/or Bayesian networks [16, 17], that have included stochastic parameters [18] and others. Nevertheless, the model of GRN dynamics that has received the maximum attention is the discrete Boolean network model, introduced by Stuart Kauffman [19]. In this model, gene expression has a binary quantification (on/off) and its temporal change in gene activity occurs in discrete-fixed steps. Besides, the state of expression of each gene is functionally related to the state of expression of some other genes (regulators), using logical rules. Even if continuous models are more realistic, they have many free parameters which are hard to constrain from experimental data, while discrete Boolean models can greatly reduce the number of these parameters while still capturing the essential network dynamics. Recent studies support that the supposedly simplistic discrete Boolean formalism, which emphasizes qualitative details, may answer realistic biological questions through experimental observations [20] and/or experimental data [21–23]. Therefore, experimentally grounded discrete Boolean GRNs models have become an important modeling tool, widely used nowadays by systems biology practitioners.

Despite the progress in understanding and uncovering such relatively low-dimensional Boolean GRN modules, we still lack analyses adjusted to unravel how such predominantly conserved regulatory modules interact with signaling mechanisms or micro-environmental cues, and also to further reveal the modular nature of the larger networks involved. We will understand the overall regulatory functionality of certain Boolean GRN results, from both the interconnection of its constitutive modules and the responses from exogenous *stimuli*. Conditioned by their topological connectivity, such exogenous inputs could alter the Boolean dynamics and consequently, in some cases, the state-space is completely restructured, *i*.*e*, state trajectories emerge when the network interacts with its entourage. Specially, studies have mainly focused on attractor transitions, because they may constitute changes in cell-fate decisions. From a Control Theory perspective, roughly speaking, the possibility of transfer the zero state or initial condition (*x*_0_) of the given system to any other state or final condition (*x*_*d*_) by using a set of admissible controls (*u*) in finite time (*t*), is known as controllability from the origin [24, 25] (or reachability). It is also said that *x*_*d*_ is reachable from *x*_0_ in finite time *t* when performed by *u*. Hence, this property captures in functional terms the structural constraints that limit the extent of manipulability where a given complex GRN can accept from its entourage [26]. In terms of structural controllability, recent published results have established a starting point to understand the role that nodes, edges, and in general topological connectivity, may be playing in the control of complex networks [27–30]. However, those results are focused on the properties of the complex networks, putting aside the nature of the system that they describe. In an attempt to contribute to such a need, in this work we propose a computer-based systematic analysis which takes advantage of the reachability property of dynamical systems, applicable to experimentally grounded Boolean GRN. Because of the inherent computational complexity that arises when studying large scale complex networks [31, 32], we restricted our study to the analysis of low-dimensional discrete Boolean GRN (at module level). Our main purpose is to gain some biologically-motivated insights about how developmental, physiological and/or environmental cues could be acting on specific genes, and consequently promoting attractor transitions.

We must point out that the here proposed controllability analysis, more properly reachability analysis, is based on a *perturbative control approach* where, in other words, the main goal is to transit from a stable or cyclic attractor to other by forcing the state variables of certain nodes in a specific time. Our method has many desirable characteristics, for instance, the node interventions guarantee to drive an initial state to the target attractor in a long term successfully, meaning that the manipulation of the node only is required during the transition of the complex network, going back to an autonomous dynamic system at the end. Moreover, if transition is available by a manipulation of a specific gene, we can directly relate its influence in cell fate transitions and the plasticity of gene regulatory networks. In addition, this approach does not require many parameters to be fixed, since it harnesses the structural controllability of the network. However, there exists other non-perturbative methods, for example, those control approaches that include feedback vortex sets [33], stable motifs [34], algebraic methods [35], and canalization-based control [36], which are also often used in biology experiments. Both non-perturbative and perturbative methods could be complementary. First approaches could help us to identify the potential nodes and then the reachability analysis could give us the suitable on-off sequence to how manipulate them. It is important to indicate that those on-off sequences are not unique, so we could implement our procedure with an optimal control approach using a Markov decision process to determine a control policy that will indicate which control to use from a given initial state to a new attractor [37].

One of the most studied GRN developmental module that has been used to uncover important aspects of early flower development, is the *Arabidopsis thaliana* floral organ specification (FOS-GRN), which comprises a useful system to address the study of structural control of complex GRN underlying cell differentiation. The FOS-GRN model in the latest version [2], successfully modeled from experimental data for the 13 genes and their interactions, proposes a robust functional module that recovers the gene activation configurations characterizing floral organ primordia (FOP) [1]. Given an initial condition, this autonomous dynamical system, where the trajectories only depend on its initial conditions, can only converge to one of ten fixed-point attractors; four of them (named *I*_*i*_, *i* = 1, 2, 3, 4) are specifically related to both the structural configuration and the dynamical behavior of the Inflorescence Meristem (IM), found in the apex of mature plant. The other six attractors correspond to sepal (SE), petal (PE1 and PE2), stamen (ST1 and ST2) and carpel (CAR) primordial cells within the Flower Meristem (FM). In consequence, FOS-GRN constitutes a model-of-choice to elucidate the role of specific genes involved in IM-to-IM, IM-to-FOP and FOP-to-FOP transitions, in the context of morphogenetic patterns of development. For example, it has been shown that a sustained modification of gen characteristic decay rate [38] on the approximated continuous FOS-GRN is sufficient to promote spontaneous attractor transitions and consequently a cell identity change. This modeling framework uncovered some of the unknown deterministic inducers that are implicated in IM-to-FOP transitions, previously hypothesized as a result of a stochastic exploration of the Boolean FOS-GRN [39]. But, three natural questions arise: *is the span of these attractor transitions expanded by the combinatorial temporal switched-on and off inputs?*

If so, *can these suitable Boolean inputs provide more solid insights about the interaction of the network with its entourage?* And even more interesting, *can they be correlated with biological observables?*. These questions are intended to be answered along this contribution, using as a case of study the aforementioned FOS-GRN module.

To set the stage, we first explore reachability properties of the Boolean network in its algebraic form [40], once a Boolean control input is connected to a specific gene. We named it reachability analysis. We then characterize the reachable subsets associated to the ten fixed-point attractors, also referred to as available trajectories. According to our results, some developmental and homeotic trajectories are tested in order to discuss the potentiality of the concerned gene in the context of biological data. To go further, we also approximate the discrete Boolean network to a continuous model and we were able to recover quantitative information concerning the influence of control inputs on the decay/saturation rates of the genes.

## Methods

### Boolean Networks Dynamics

A Boolean Network (BN) is a discrete-time dynamical system of *n* Boolean variables *x*_*i*_ ∈ Δ_2_ = {1, 0}. The state of expression of the BN at time *t* is given by a vector of *n* Boolean variables [*x*_1_(*t*)*x*_2_(*t*), …, *x*_*n*_(*t*)]^*T*^. Hence, *x*_*i*_ ∈ Δ_2_ is the state of expression at time *t* of the *i*-th gene, and changes over time according to the dynamic equation:

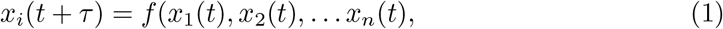

where {*x*_1_(*t*), *x*_2_(*t*), … *x*_*n*_(*t*)} are the regulators of the gene *x*_*i*_, *i* = 1, 2, …, *n*.

*f*_*i*_ : *D*^*n*^ → *D* is the Boolean function (or logical function) related to gene *x*_*i*_, constructed in agreement to the combinatorial action of the regulators. *τ* is the necessary time taken by a gene to change the state of expression of its regulators, fixed in *τ* = 1. Hereafter, we suggest to see Supplementary Section 1 for further information about Semi-Tensor Product and notation.

### From a BN to a discrete-time dynamical system

To convert the dynamics of the BN described by Equation 1 into an equivalent algebraic (*i*.*e*., in terms of a conventional discrete-time linear system), we define 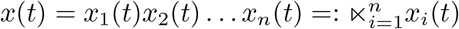. Moreover, there exist structure matrices [40], 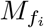, *i* = 1, 2, …, *n*, for every logical function *f*_*i*_ associated to 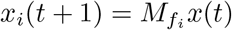, *i* = 1, 2, …, *n*. In addition, 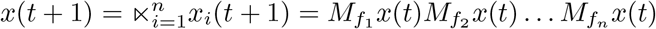. Hence, *x*(*t* + 1) can be expressed as:

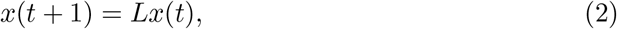

where 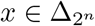, and *L* is called the transition matrix of BN. We must point out that Equation 2 is enough to describe the full dynamics of the concerned GRN.

Once the discrete-time Boolean GRN is transformed and expressed by Equation 2, the attractors of the network can be defined as follows [40]: if we consider a state 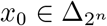, it is called a fixed-point attractor if *Lx*_0_ = *x*_0_, while we say that {*x*_0_, *Lx*^0^, … *L*^*k*^*x*_0_} is a cyclic attractor with length *k* if *L*^*k*^*x*_0_ = *x*_0_.

### From a BN to a Boolean control network

To characterize the reachable subsets associated to the previous identified attractors, we first transform a BN described by Eq. 2 into a Boolean Control Network (BCN) under three assumptions: (1) the network is synchronous, (2) each node can be manipulated at any time, and (3) there is no restriction about the number of control inputs. Those control inputs or controllers (used indistinctly) have also a Boolean behavior, more precisely called Boolean sequence, and they are represented by a variable u. Here we adopted the next system of equations:

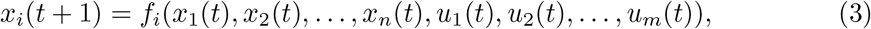

where *x*_1_(*t*), *x*_2_(*t*), …, *x*_*n*_(*t*), *u*_1_(*t*), *u*_2_(*t*), …, *u*_*m*_(*t*) are the regulators of the gene *x*_*i*_,*i* = 1, 2, …, *n*, which now also depend on the *m* control inputs 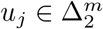, *j* = 1, 2, …, *m*. *f*_*i*_ : *D*^*n*+*m*)^ → *D* is the Boolean function related to gene *x*_*i*_.

### From a BNC to a discrete-time bilinear dynamical system

The same procedure to obtain the algebraic form of BN holds for the BCN described by Equation 3, but in this case we have a bilinear system expression since, if we define 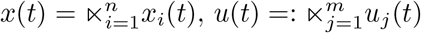 and multiplying all together yields:

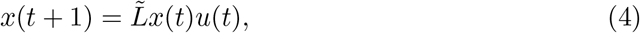

where 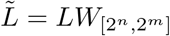. We can see that for time *t* = 0, 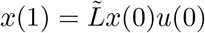, for 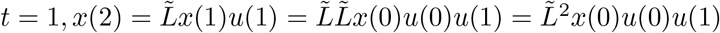, then for *t* = 1, 2, …, 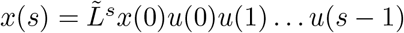. Thus,

#### Theorem 1

**([41])** *x*_*d*_ *is reachable from x*_0_ *at the s-th step by controls of Boolean sequences of length s if and only if:*

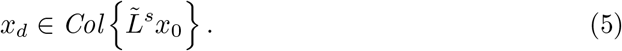

In cases where Equation 5 holds, then the control are defined by:

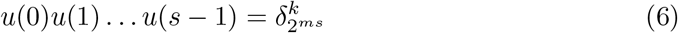

### Reachability analysis

To find the potential nodes that promote trajectories between attractors in a Boolean GRN, we propose the Algorithm 1 based on the result given by Equation 5. Basically, once the attractors are identified, we test if adding a control input to the *i*-node (through either AND and OR connector) is sufficient to promote a transition from one attractor to another. If so, we say that the final attractor *x*_*d*_ is reachable from the initial attractor *x*_0_ by a control input *u* via the *i*-node. Such available trajectories promoted by the action of control inputs on the nodes can be tested by a discrete-time simulation.

### Approximation to a continuous Boolean Control GRN model

It has been shown above that Boolean GRN models transformed into its corresponding equivalent algebraic forms, are sufficient to characterize developmental trajectories in terms of reachable subsets, and consequently elucidate the potentiality of the manipulated genes to promote these transitions. However, discrete approaches do not contemplate aspects such as the differences in genetic expressions decay rates, saturations rates and other quantitative parameters of grounded on data biological GRN. Thus, it becomes necessary to approximate such Boolean models to continuous systems, in fact, previously approaches have been used to this end [2, 3].

#### Algorithm 1

Characterization of the reachable subsets associated to the fixed–point attractors in a Boolean GRN

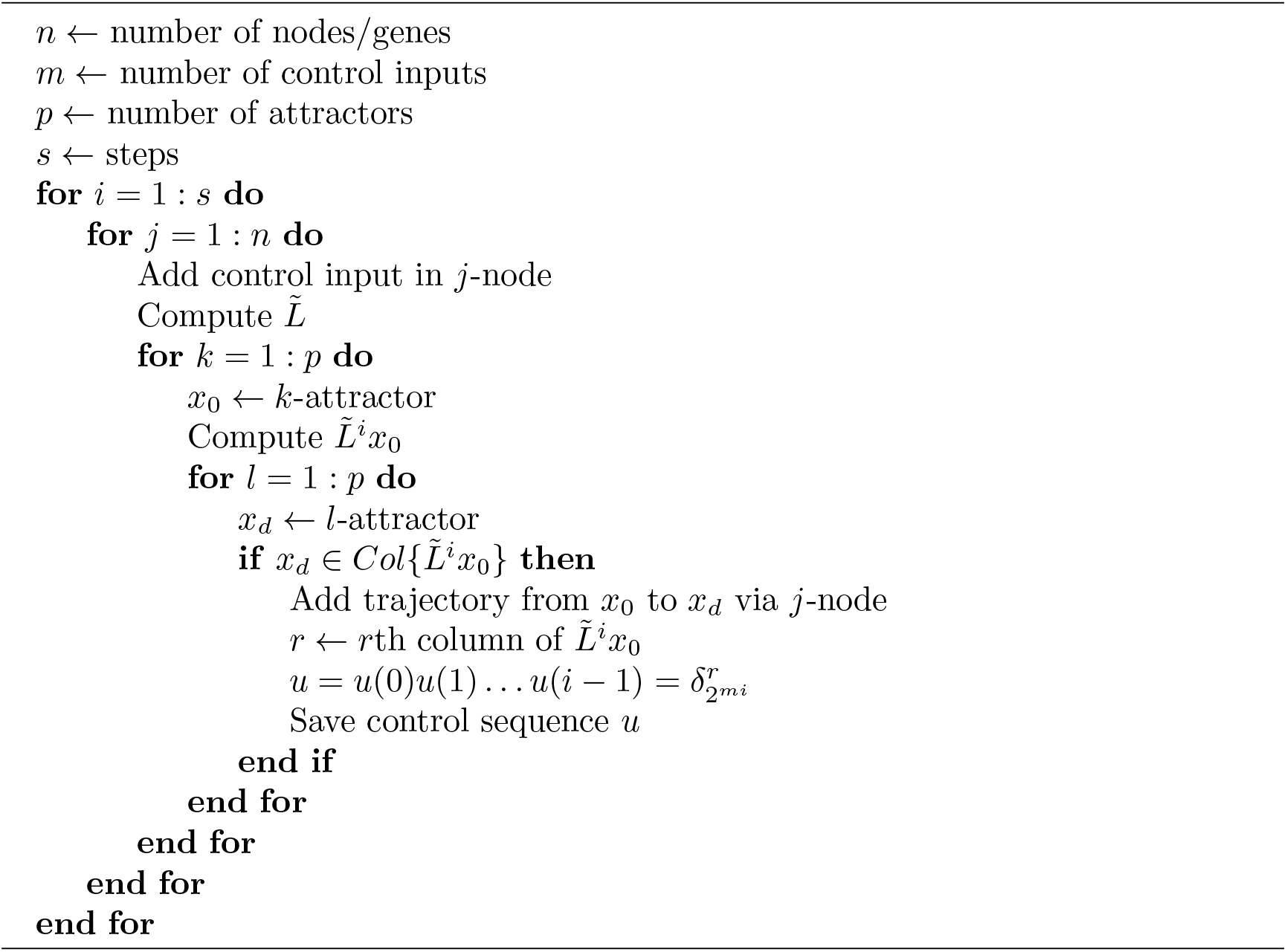

Particularly, we are interested in testing the Boolean control inputs obtained by Algorithm 1 on the continuous GRN and recovering the previous characterized developmental trajectories on the Boolean GRN. Therefore, we first select a trajectory from one attractor to another and proceed to add the control input on the Boolean GRN, then, we transform each Boolean 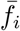, which depends on the regulators and *u*, into a continuous 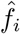 as follows:

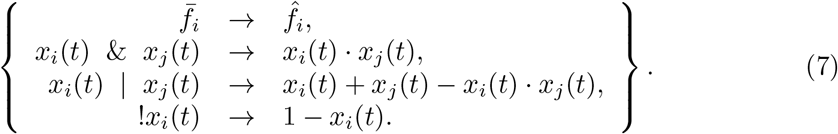

Afterwards, we adopted a system of Ordinary Differential Equations (ODE’s) of the form:

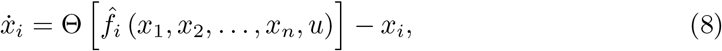

where *u* is a time-dependent parameter, which is interpolated to obtain the value of time-dependent terms at a specified time (*i*.*e*. 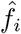). Then, we consider that the input function displays a saturation behavior characterized by the following logistic function:

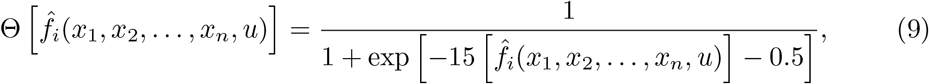

where *ϵ* is a threshold level (usually *ϵ* = 0.5), and *b* stands for the saturation rate. We notice that for *b >>* 1, the input function displays a dichotomic behavior. The value of the parameters *b* is chosen as the smallest integer value able to recover the same fixed-point attractors and their basins of the Boolean model. Finally, the numerical solution of the ODE’s system is conducted using a standard numerical integration solver available in Matlab™.

## Results

### Analysis framework

The proposed reachability analysis of a Boolean GRN, which can give us insights into the role that the individual genes have in the context of the network as a whole, consists of six main steps, schematically depicted in Figure 1.

**Fig 1.**
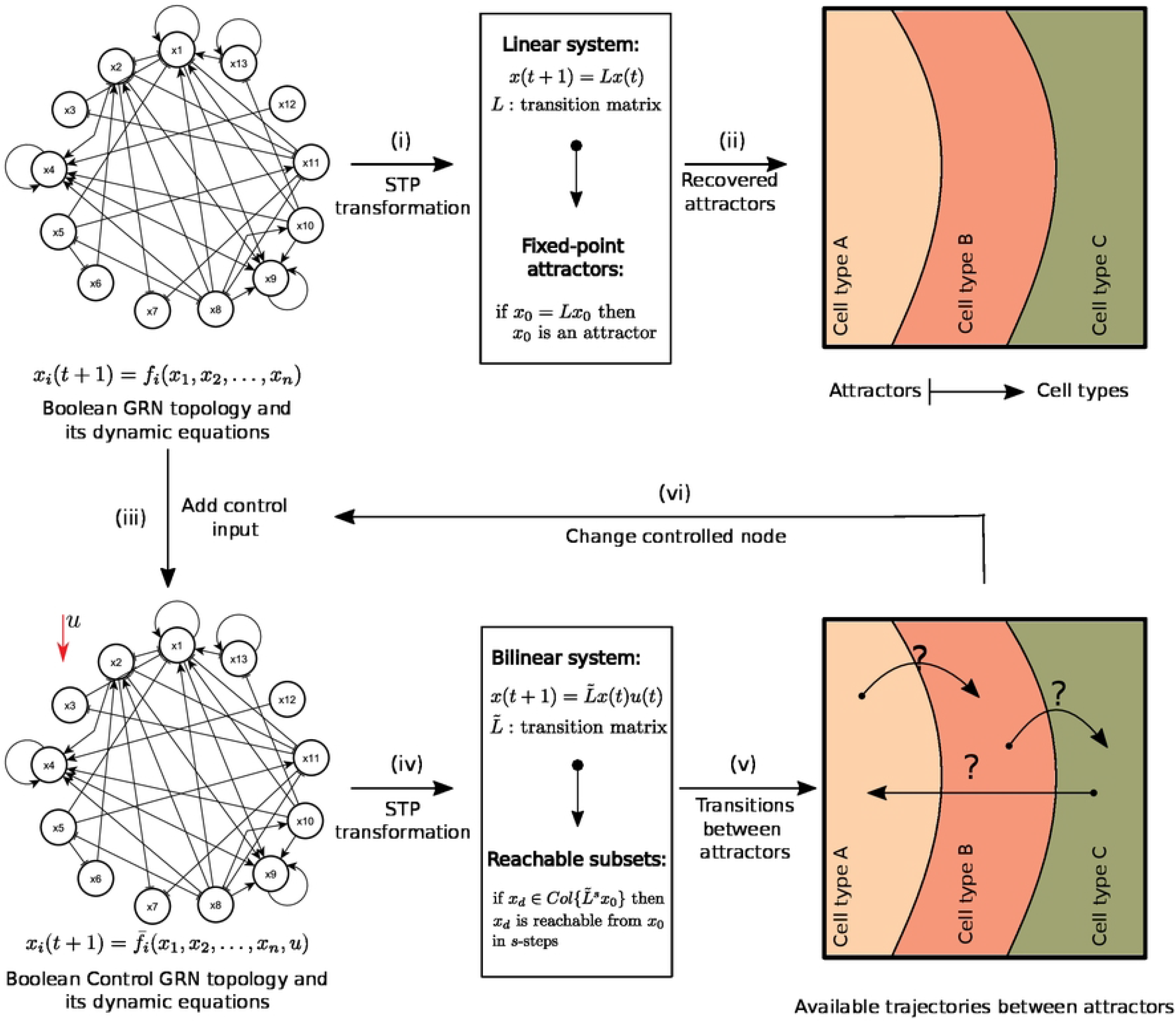
Overview of the reachability analysis of a Boolean GRN. (i) The starting point is an experimentally grounded low-dimensional GRN. Its set of logical rules are rewritten into algebraic operators via semi-tensor product, therefore, the Boolean GRN is transformed into a discrete-time linear system. (ii) The attractors (fixed-point or cyclic) are identified in the linear system, after an exhaustive exploration for every initial condition. (iii) A Boolean control input is connected to a gene, restructuring its corresponding logical rule. (iv) Thus, the new Boolean Control GRN is transformed into a discrete-time bilinear system, as stated above. (v) If the Boolean control input is able to promote a transition from an initial attractor to another, we say that this is an available trajectory. Finally, (vi) the process is repeated until all the genes are explored.

### Dynamic analysis of FOS-GRN and its transition matrix

The most recent version of *Arabidopsis thaliana* flower organ identity GRN (conformed by 13 genes and referred to FOS-GRN) reported in [2], is used here as a case-of-study.

FOS-GRN is described as an autonomous dynamical system in Boolean terms (see the set of Boolean functions on Supplementary Table S1) and characterized from experimental evidence. We transformed the FOS-GRN, applying the STP approach into the discrete-time linear system given by:

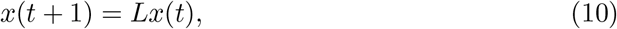

where *L* is an integer matrix of dimensions 2^13^ *×* 2^13^ (*n* stands for the number of genes in the network), also known as the transition matrix of the network (see Materials and Methods). Then for all 2^13^ possible initial conditions *x*_0_, we identify the attractors or stable configurations of the activity of the network (corresponding to specific gene expression profiles), *i*.*e*. all solutions to the equality *Lx*_0_ = *x*_0_. This network recovered ten fixed-point attractors as we expected, and its gene activation configurations are presented in Supplementary Figure S1. Four of them are the ones associated with both the structural configuration and the dynamical behavior of the inflorescence meristem (IM) found in the apex of mature attractors corresponding to sepal (SE), petal (PE1 and PE2 except for the state of activation of the gen UFO), stamen (ST1 and ST2, the same case as petal) and carpel (CAR) primordial cells, which emerge from Flower Meristems (FM’s) formed in a helicoidal pattern from the flanks of IM [1, 21]. The basin size of each attractor, a result derived from the analysis of the transition matrix, might be important since it can be related with if an attractor transition is more likely to be done. Thus, the basin size of *I*_1_, *I*_2_, *I*_3_, *I*_4_, SE, PE1, PE2, ST1, ST2, and CAR attractors is 136, 136, 72, 72, 812, 12, 824, 94, 3064 and 2970 states, respectively. As we expected, the total sum is just the number of all possible gene expression profiles. In Figure 2, we show each attractor and their spatial position on meristems, as well as a unique tag for each gene profile *D*_*e*_ (decimal equivalent of its gene profile in binary code).

**Fig 2.**
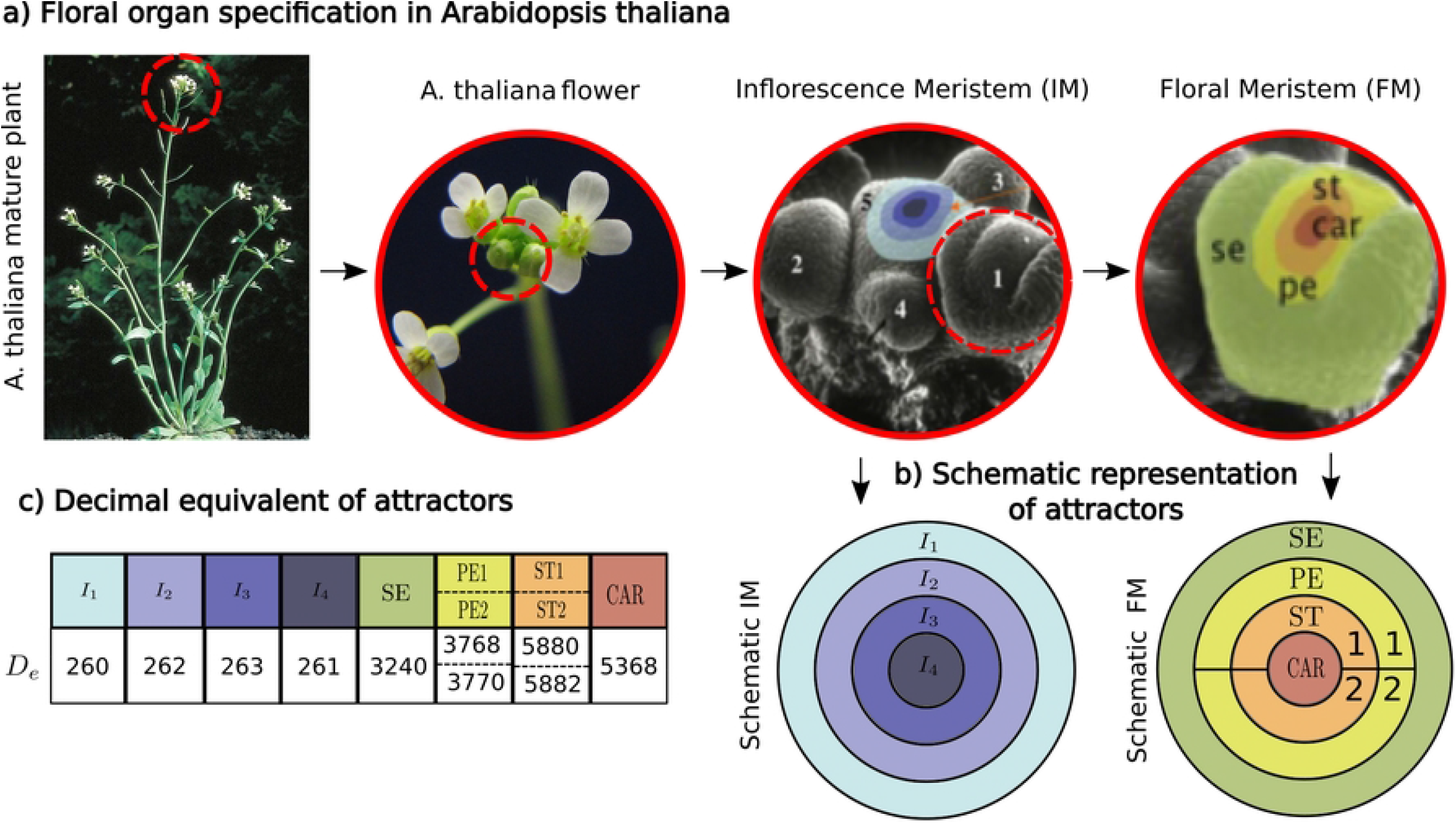
The ten fixed-point attractors of the FOS-GRN. a) Four attractors are associated to the inflorescence meristem (IM), where we can distinguish four sub-zones (*I*_*i*_,*i* = 1, 2, 3, 4). Flower meristems (FM’s) arise from flanks of the IM (1, oldest; 5, youngest). The other six attractors correspond to this FM, which is divided into four concentric regions that will eventually give rise to the flower organ primordia: sepals (SE), petals (PE), stamens (ST) and carpels (CAR). b) For illustrative purposes on further sections, we represent these attractors with colored concentric circles. c) To identify all the attractors, we converted the binary code of the gene profile to a decimal equivalent (*D*_*e*_). In cases such as PE and ST, gene UFO can be expressed or non-expressed, so two different *D*_*e*_ can be assigned.

### Reachable subsets of the FOS-GRN

From a Control Theory perspective, the possibility to transfer the given zero state or initial condition *x*_0_ to any other state or final condition *x*_*d*_ under a suitable control input *u* in *finite* time *t* is called as *controllability from the origin* or, more often, reachability, which is a structural property of the dynamic system [42]. In fact, this statement agrees with our goal to expose the potential genes (hereafter referred to *portiers*) of the FOS-GRN, which are involved in attractor transitions, since reachability can be extended to Boolean systems, considering a finite number of steps *s* as a finite time. So exhaustive characterization of the reachable subsets (also known as *reachability analysis*) associated to the fixed-point attractors as *x*_0_, is sufficient to elucidate the portiers of the FOS-GRN. Note that, however, we do not establish the molecular nature of the control elements *u*, but we later provide some examples, discussing how the entourage could be interacting with the FOS-GRN module in accordance with available biological data.

In order to explore such *portiers*, we transformed the FOS-GRN into a Boolean Control Network (BCN), adding a control input on the *i*-th gene. We tested separately two possible Boolean operators (as connectors) between original Boolean functions *f*_*i*_ and *u* to create the new Boolean function 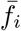 the OR operator (switched-on), and the AND operator (switched-off). Thus, we adopted a new system of Boolean equations:

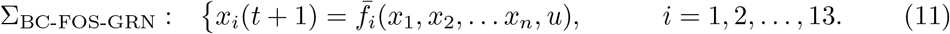

These equations now also depend on *u*. Afterwards, we applied the STP transformation but now for the BCN, and the system became a discrete-time bilinear system of the form:

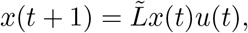

where 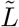 is a matrix of dimensions 2^14^ *×* 2^13^, also called as a transition matrix of the BCN (see Materials and Methods). Then, to characterize the reachable subsets of the FOS-GRN given the set of attractors and the computed 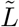 we performed the Algorithm 1 proposed in the Methods section for *s* = 1, 2, …, 10.

### Portiers of the FOS-GRN

As we mentioned above, we called *portiers* those potential genes, characterized as a result of the reachability analysis, so that by changing their level of expression through the interaction with the control input *u*, promote a trajectory from a given attractor to another given attractor. Particularly for our case study, a total number of 79 available trajectories were found on FOS-GRN -at least under the two connectors tested here- and they are summarized in Table 1 (for more details about *s* and *u*, see Supplementary Table S3 and S4). Interestingly, we noticed for the first time that some trajectories between attractors are not allowed at all. For example, a considerable gap appeared on those transitions from floral primordia to inflorescence-like meristems, indicating that, it is difficult to return to a previous stage once the developmental program of flowering initiates. So that mutants that produce flower with inflorescence-like characteristics, for example *ag-1* mutants grown in short day conditions [43], are probably not caused by a return to an inflorescence state but they may now they have a novel potential of the IM state to choose between two cell-fate decisions: floral or inflorescence-like meristems, as has been shown for other MADS mutants by the epigenetic landscape analysis of the XAL2 regulatory network module [44]. Furthermore, note that another gap appears on transitions from inflorescence-like meristems to PE1 and ST1 attractors, we suppose it is a consequence of their basin size (12 and 94 states, respectively) in contrast to the PE2 and ST2 (824 and 3064 states, respectively).

**Table 1.**
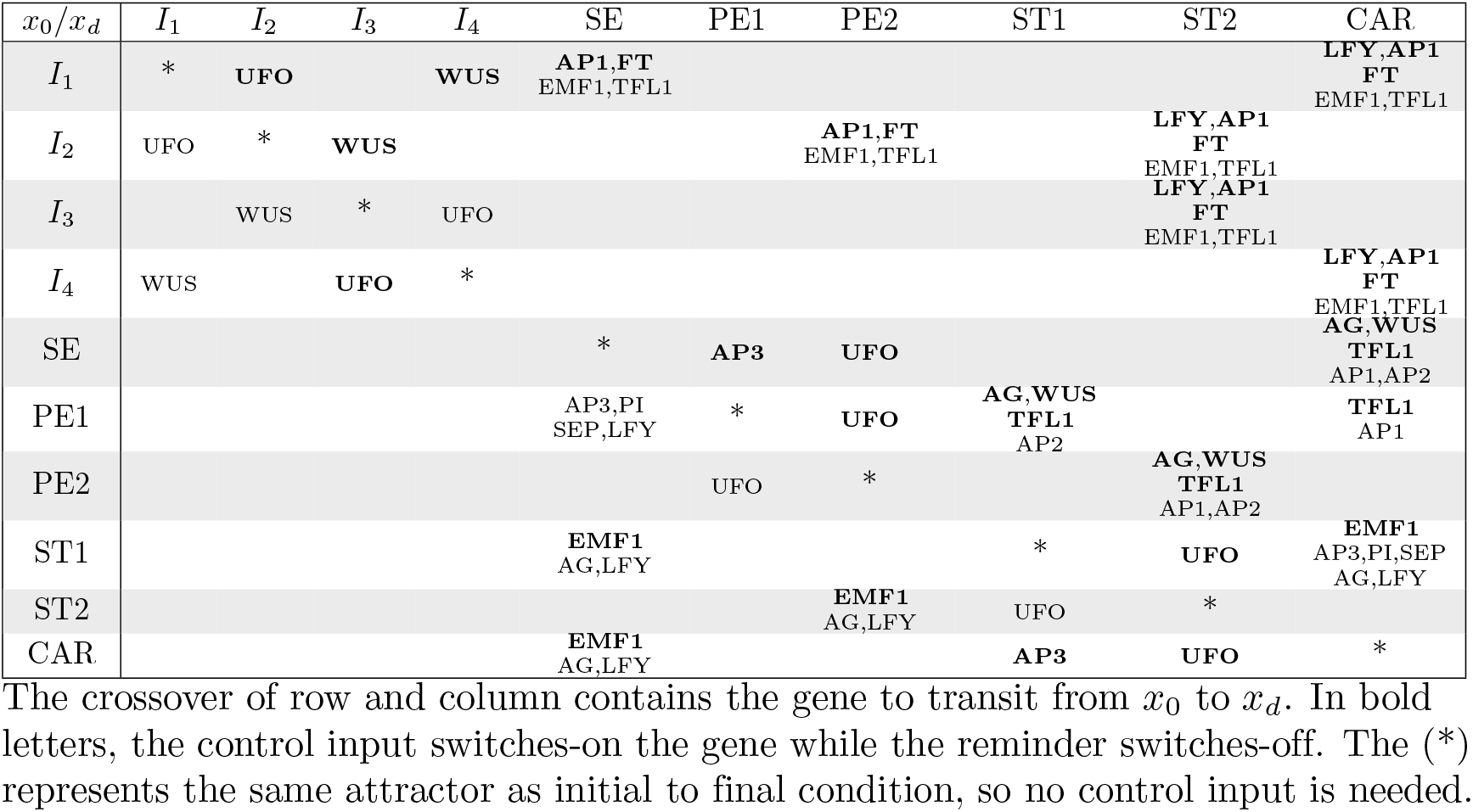
Available trajectories among all possible pairs of attractors induced by potential genes.

On the other hand, we can interpret those available trajectories either as (1) developmental or (2) homeotic. First, we focused on the trajectories that occur in normal development, for instance, inside of the sub-zones of IM and from IM to floral primordia. Both UFO and WUS are necessary for the transitions of the sub-zones (*I*_*i*_) of the IM, as a consequence of changes in the positional information caused by the displacement of the cells since the shoot apical meristem is constantly growing. We certainly found the developmental trajectory:

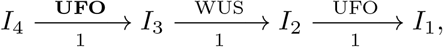

meaning that (in a compact notation), starting from *I*_4_ attractor and switching-on UFO (in bold letters), *I*_3_ will be eventually reached, thus being on *I*_3_ and switching-off WUS, *I*_2_ will be now reached; and finally, starting from *I*_2_ attractor and switching-off UFO, *I*_1_ will be the final attractor, i.e. the final point of the corresponding morphogenetic trajectory. To illustrate these induced trajectories, we generated a graph with two sections; at the top we plotted the initial attractor state and its progressive change under the effect of the controllers, and at the bottom the level for each controller.

Figure 3.a) shows the graph of the transitions inside IM (from the youngest to the oldest sub-zone), red line indicates a switch-on, whereas the blue line shows a switch-off. We see that the attractors (tagged with *D*_*e*_) are visited, and such transitions only require one step to be induced, consistent with the necessary steps presented in Supplementary Table S3.

**Fig 3.**
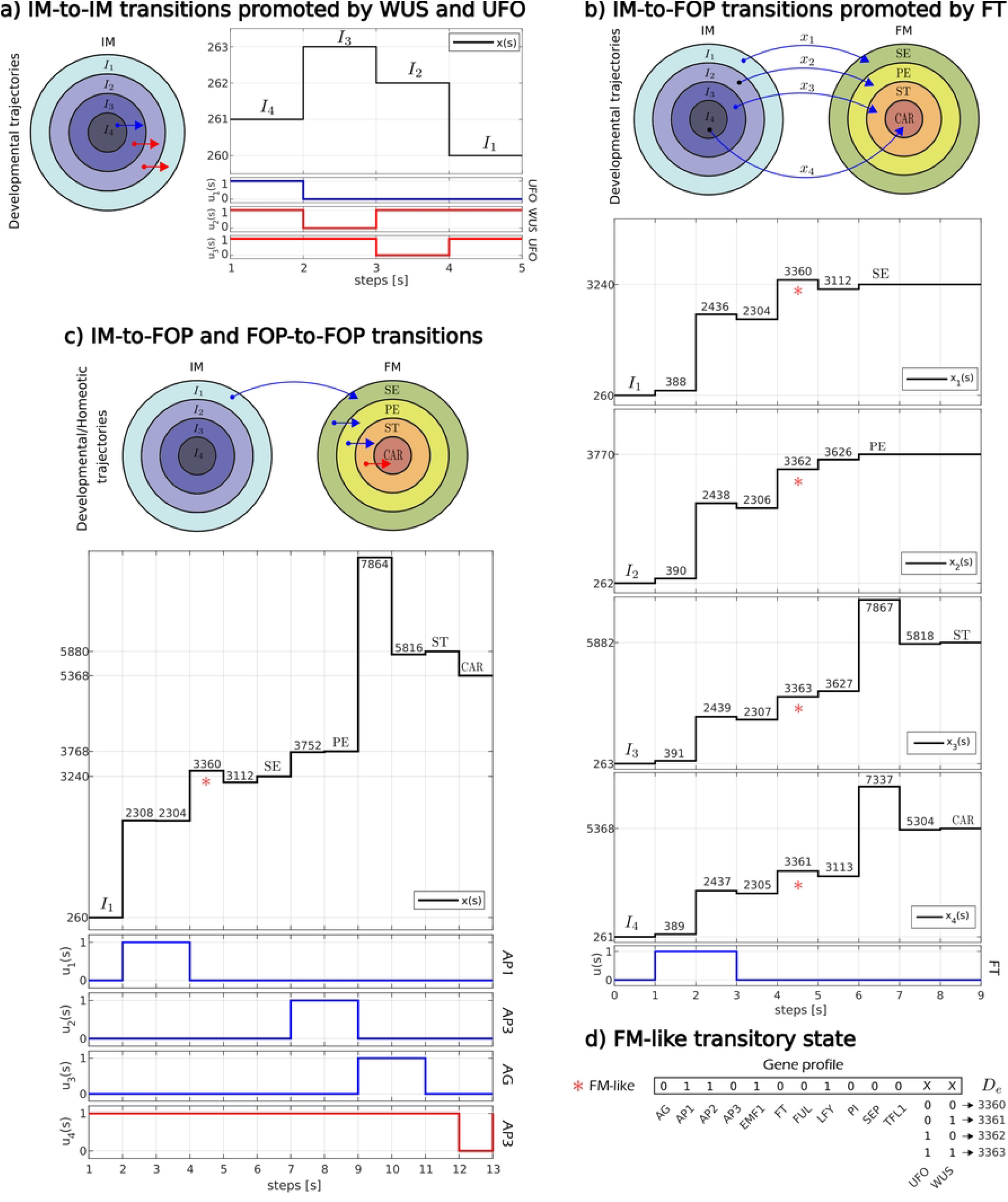
Developmental/Homeotic trajectories on the IM and FM under control inputs on *portiers*. a) A trajectory from the youngest (*I*_4_) to the oldest (*I*_1_) sub-zones of the IM is tested by suitable manipulation of WUS and UFO. The red line indicates that the controlled gene is switched-on while the blue lines indicate that the genes are switched-off. b) Four developmental trajectories *I*_1_-**FT**→SE, *I*_2_-**FT**→PE2, *I*_3_-**FT**→ST2, and *I*_4_-**FT**→CAR, are also tested. Interestingly, a suitable manipulation of the *portier* FT promotes all these attractor transitions. In addition, we added a FM-like tag to those transitory states that correspond to the configuration of the FM before reaching the desired attractor and the gene profile that holds for all the cases is shown at the bottom in d). A combination of developmental and homeotic trajectory is shown on c). Here we mimic the spatial-temporal sequence of floral organ specification in real plants.

Eventually, such IM’s will give rise to FM’s, and they will sub-differentiate in the flower organ primordia. It has been reported in [44–46], that some of the characterized regulators which determine these meristematic cells are LFY, AP1 and AP2 while, on the other hand, the activity of the FM is counteracted by the gene TFL1 [47].

Consequently, an expected attractor for floral meristems will include on its gene profile a LFY=AP1=AP2=1 and TFL1=0, since UFO and WUS are only involved on the transitions of the sub-zones of the IM. Although the FOS-GRN do not recover an attractor with the configuration of the floral meristem, we reasoned that the induced trajectories (by *portiers*) from the inflorescence meristem to the floral organs may be passing by a transitory state that corresponds to the FM. To test this, we selected trajectories from:

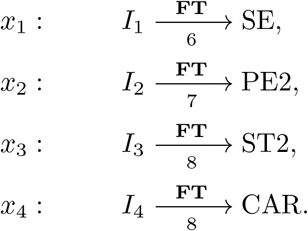

It is important to point out that the manipulation of FT on this testing is onñy chosen for illustrative purposes. Indeed, LFY, AP1, TFL1, and EMF1, could also work. Figure 3.b) shows how each of the four cases of induced trajectories (*i*.*e. x*_1_, *x*_2_, *x*_3_, and *x*_4_) have a transitory state where AP1, AP2, LFY and TFL1 have the expected configuration for FM-like attractor but only in a transitory form (for further information about the these states, see Supplementary Figure S3). Besides, according to Table 1, the *portiers* AP1, LFY, EMF1 and TFL1 also work. While FT, AP1 and LFY must be switched-on, EMF1 and TFL1 must be switched-off. Indeed, when these nodes are manipulated the FM-like transitory state is also visited. This is actually what happens during development, for example, long day grown plants, where FT is active, produced fewer secondary inflorescences than short day grown plants where FT is not transcribed [48]. But it is even more interesting that floral homeotic genes such as AP3, PI and AG, despite being necessary, are not enough to form floral organs, in agreement with our previous hypothesis on the ABC model of floral patterning [21]. Actually, the control of AP1 in the 35S::AP1-GR ap1 cal plants that grow inflorescence meristems until the modification to floral meristems when treated with dexamethasone, showed that only one treatment of dexamethasone is enough to form the four types of floral organs [49]. These experiments were done after the publication of the FOS-GRN [21]. Therefore, the reachability analysis can make predictions on how biological modules may be controlled.

Secondly, we focus on the trajectories that can be explained as homeosis, which are associated to transitions between floral primordial (FOP). Unlike IM-to-FOP developmental trajectories induced by *portiers* LFY, AP1, FT, EMF1 and TFL1, FOP-to-FOP homeostatic trajectories can be promoted by *portiers* AP2, AP3, PI, AG and SEP, which are recognized by their mutant homeostasis flower phenotypes. Even WUS and UFO are able to cause such transitions, showing that alterations in floral patterning can be caused by changing spatial cues in the meristem. Now, we take as an example of a control scheme a node not presented in the network, the BELLRINGER gene [50]. In the blr-4 missense mutant AG is depressed, and some flowers develop carpels in place of sepals in the first whorl. The whorl 1 and carpels are often fused and form a gynoecium that encloses the rest of the flower [51]. In fact, such switch-on of AG to transit from sepal to carpel primordium is evidenced by our proposed reachability analysis.

### The role of the *portiers* is maintained on the continuouos FOS-GRN model

Until this point, we were only interested in qualitative changes on the dynamics of the FOS-GRN under the effect of Boolean sequences connected to *portiers*. However, quantitative information as the minimum time of manipulation of these potential genes and their impact on the relative decay/saturation rates of the rest of the genes, have not yet been explored. To illustrate this, we first selected a composed developmental/homeotic trajectory:

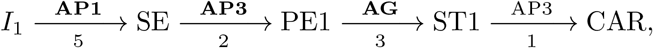

which mimics the sequence of attractor transition previously recovered by different levels of noise in the updating rules of the network [38]. We then tested the control inputs obtained through our proposed reachability analysis on the Boolean FOS-GRN (see Supplementary Figure S3). We also approximated the Boolean dynamic of the FOS-GRN without control inputs following a Glass piece-wise continuous approximation (see Methods). We tested a range of values of *b* = *i* for *i* = 1, 2, …, 30. For the input-state response, we noticed that the parameter *b* ≥ 20 is able to recover the same attractors and basin size of the Boolean network. On the other hand, the *ϵ* threshold were fixed in 0.5 for simplicity (the set of continuous functions are presented in Supplementary Table S4).

In order to find the minimum time of manipulation of the *portiers* by the control inputs *u*_*i*_ in the Boolean Control FOS-GRN, we separated the composed developmental/homeostatic trajectory into four individual transitions, and consequently four different ODEs systems were also approximated as stated above. For each attractor transition, we took the initial attractor as a *x*_0_ of the ODEs system, and we then increased the duration of the *j*-th control input (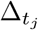), interpolating its value to obtain the value of the time-dependent terms at a specified time. In this way, a plot is generated with the parameter value of 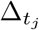 in the *x*-axis, whereas in the *y*-axis the total sum of the single gene expression values for the *n* genes is specified. Figure 4.a) shows the bifurcations diagrams obtained for each attractor transition, and as we can see, despite the *portier* AP3 promotes the trajectories from SE to PE1 and from ST1 to CAR, the necessary 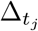 is different. Moreover, we considered an extra time for each interval 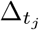 in such a way that the *portier* and the rest of the genes have enough time to relax to its new state after the manipulation by the control input (see Supplementary Figure S4). We finally integrated all the individual transitions in Figure 4.b). In the axis we show the relative concentration of the 13 genes involved in the FOS-GRN, and in the *y*-axis is included the time intervals for the four control inputs. This plot suggests that there exists a *minimum trigger threshold* which, once it has been crossed, provokes an imminent attractor transition.

**Fig 4.**
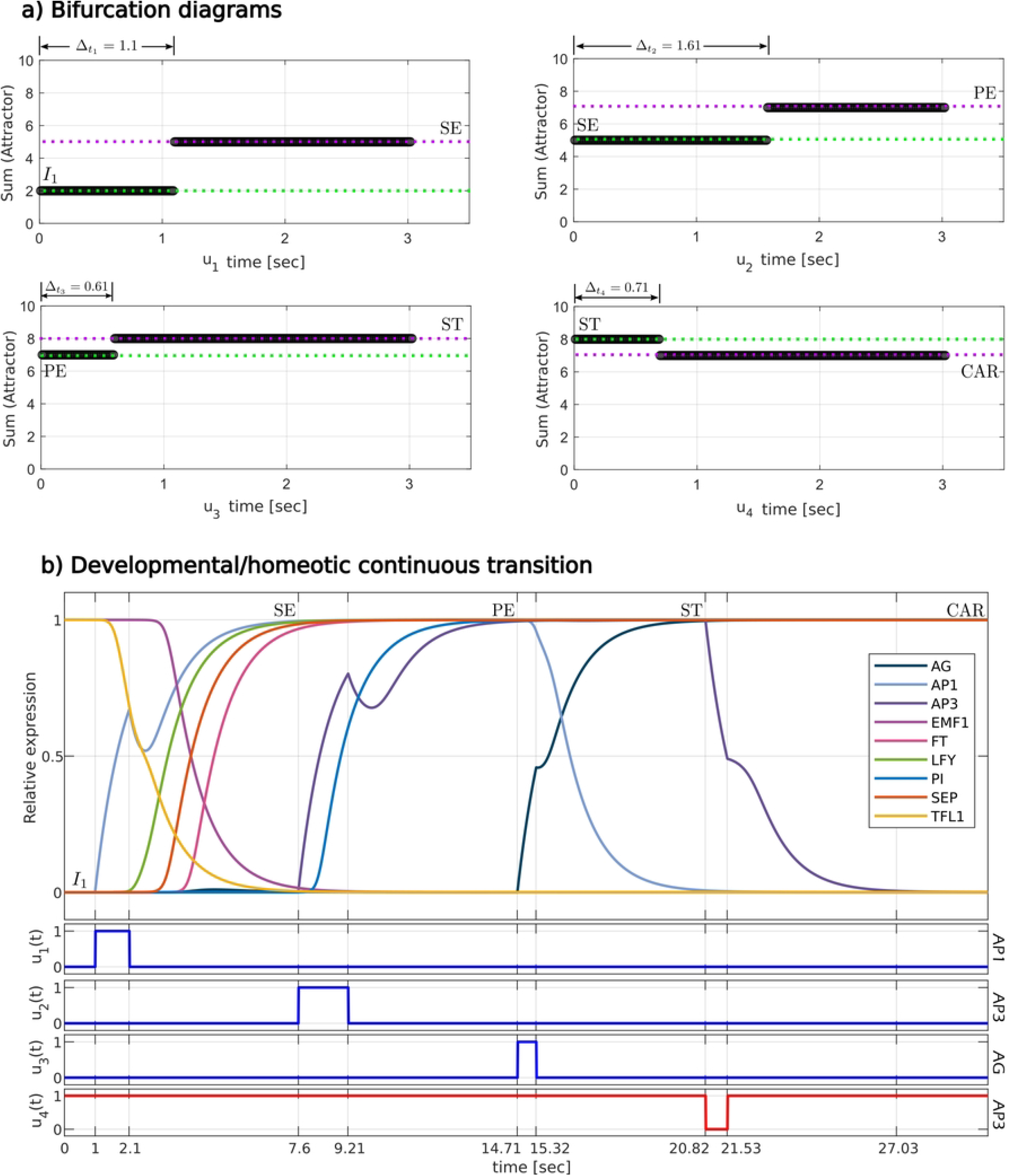
A developmental/homeotic trajectory in the FOS-GRN that mimics the sequence of gene profiles observed in real flowers. The trajectory *I*_1_–**AP1**→SE–**AP3**→PE1–**AG**→ST1–AP3→CAR is tested. First, in a) we show the bifurcation diagrams for the minimum time of manipulation of each gene by the control input *u*, sufficiently enough to produce qualitative changes in the sum of the active genes. Based on these times, each control input is built and their values are interpolated in the solution of the ODE’s system. In b) we show a continuous plot with the relative concentration of each gene under the computed control sequences.

## Discussion

A quintessential issue in Control Theory is to find a control input which drives a dynamic system from an initial to a desired state in finite time, *when possible*. In fact, there exist several ways to find such control input depending on the specific nature of the concerned dynamic system. For example, in some multi-stable systems, suitable variations on a single parameter are sufficient to change from one steady-state (or attractor) to another. In this same direction, such transitions in experimental grounded Gene Regulatory Networks (GRNs), promoted by an external stimulus in a particular gene, originated from responses of changes in physiological, environmental and/or developmental cues that could implicate a change in cell-fate decision during development. Some authors have proposed different methodologies to capture the effect of changes on the level of expression of the genes by external elements in GRNs that underlie realistic cases of cell differentiation. This is the case of the bifurcation analysis of a single parameter [20, 38], the stochastic combinatorial Boolean interventions [57], the uncertainty in the updating Boolean rules [39], etc., but fewer have used mathematical approaches based on underlying structural properties given by their topology and their dynamics.

Here, we propose a computer-based analytical procedure which harnesses the structural reachability property of low-dimensional synchronous Boolean GRN. As a concrete study case, we use the regulatory module underlying floral organ determination in *Arabidopsis thaliana* during early stages of flower development (FOS-GRN), because it constitutes an ideal model-of-choice for the study of differentiation and morphogenesis in multicellular organisms [38, 39]. Our proposal is built around the Semi-Tensor Product approach [40] (STP), an algebraic mathematical tool that enables us to convert the Boolean FOS-GRN, a discrete-time dynamical system, into a conventional discrete-time linear system (or algebraic form), through the rewriting of the logical rules as algebraic operators. As a result of such transformation, the transition matrix of the network *L*, which is a Boolean square matrix, is unique and enough to fully describe the network dynamics. Additionally, *L* also comprises topological structural information about the fixed-point attractors, cycles, transitory periods and basin of attractors. In this manner, we recover the ten steady-state configurations corresponding to the observed gene activation configurations (see Figure 2). Thus, all classical methods and conclusions for linear systems can be used to analyze the dynamics of the network, including control techniques, em e.g., state feedback control [28, 53]. However, proposed theoretical controllers that use these methods are quite complex to have a realistic implementation (as sophisticated control techniques that take the form of engineered biomolecular interactions), and do not provide useful insights about the role of the genes in the FOS-GRN as a whole.

As stated above, we transformed the Control Boolean FOS-GRN, that occurs as a result of connecting an external Boolean input to a specific gene, into a discrete-time bilinear system. Right away, the new network transition matrix encompasses structural reachability properties of the Boolean Control FOS-GRN. We say that a state is reachable from an initial state, if a qualitative change on the state-space configuration is observed when an external Boolean input is connected to a specific gene, modifying its state to describe knockout (Off) or overexpressed (On) states, and changing the corresponding updating rule for the gene during simulation. Despite the advances in reachability analysis of GRNs [54], the exploration of the reachable subsets corresponding to attractors in experimental grounded GRN, that explains the role of the genes on such attractor transitions, has been left behind. So, we focused exclusively on attractor transitions and thus we limit the scope of our conclusions.

We found that there exist 79 different attractor transitions (see Table 1) promoted by potential genes or *portiers*, in the Boolean FOS-GRN, and they are consistent and complementary with the gene decay rates approach. Such *portiers* obtained through the reachability analysis play a role in (1) transitions among the sub-zones of the inflorescence meristem (IM-to-IM), (2) transitions from inflorescence-like attractors to flower organ primordia (IM-to-FOP), and (3) transitions among the different floral organs (FOP-to-FOP). For instance, our results in Figure 3 showed that deterministic perturbations on the gene FT, a key component of the photoperiodic pathway in Arabidopsis thaliana [48, 55], promote IM-to-FOP transitions, supporting the hypothesis that biological systems not only cope with random perturbations, but also with deterministic signals or non-random inducers [39]. In the four simulations, an unexpected transitory appears, and its gene profile correspond to a floral meristem (FM) identity. Even though FOS-GRN do not recover it, it emerges as a direct consequence of inducible IM-to-FOP transitions. Interestingly, this result enables us to speculate that reachability analysis can also give us information about intermediate states that are significant during cell-fate decisions. Furthermore, we demonstrated that such potentiality is not only conserved in the approximated continuous Boolean FOS-GRN, but also reveals the existence of a trigger threshold which is crucial in the transitioning between one attractor and another.

As has been seen, the reachability analysis allows visualizing of why certain phenotypic transitions are possible and why others are not, due to the structural restrictions to which the gene regulatory networks are subject. Consequently, the analysis of reachability, through methods such as the one we proposed here or other similar ones, can inform us about what enhances or restricts biological development. This has a particularly important significance in the context of the study of the interplay between phenotypic plasticity and evolution. In fact, the absolute robustness of the phenotypic response prevents the execution of evolutionary dynamics in response to the selective pressure imposed by the environment. The evolutionary process then favors the plasticity of biological development and such *dynamic plasticity needs to be quantified in terms of variability of the reachability of the gene regulatory networks involved*.

In a more general context, reachability analysis could also provide understanding tools of, for example, those question involved in some human diseases. Nowadays, aspects related to chronic-degenerative diseases, implicate cell dedifferentiation and cell reprogramming which result from abnormalities in the cell regulation [56]. Therefore, the elimination of health problems in this context are also subjected and necessary to control methodologies involving the design of strategies of cell reprogramming in order to return cells to a suitable state or to revoke the unhealthy trajectory [52]. We strongly believe that the reachability analysis method proposed here can be used successfully to address such kinds of problems.

Following the approach shown in this article, we are currently exploring the reachability properties of the network that underlies the epithelial-mesenchymal transition in the context of epithelial cancer [58]. One of the most remarkable advantages of the systematic treatments that characterize the study of genetic regulation by the methods of systems biology lies in the possibility of exploring broad classes of dynamic systems. The methodology presented here to clarify the interaction of some qualitative aspects of the interaction between gene regulatory networks and their environment allows, even with its limitations, to demonstrate basic principles that govern the structure and functionality of these networks. Much remains to be done in particular with regard to the study of large-scale gene regulatory networks. Approaches conceived for this purpose, such as those that make use of process hitting [59] and, in general, methods of verification of models from theoretical computing, can be complemented with what is revealed here.

For more information, see Supplementary material.

## Supporting information

**S1 Fig. The ten fixed-point attractors of the FOS-GRN**.

**S2 Fig. FM-like transitory state appears on IM-to-FOP transitions**.

**S3 Fig. Developmental/homeotic trajectory on the Boolean FOS-GRN.**

**S3 Fig. Developmental/homeotic trajectory on the Boolean FOS-GRN**. Relative concentration of genes on induced trajectories. Graphs generated from the numerical solution of the ODEs system after the exploration of 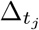.

**S1 Table. Boolean functions of the FOS-GRN**.

**S2 Table. Summary of available trajectories on the FOS-GRN Part I**.

**S3 Table. Summary of available trajectories on the FOS-GRN Part II**.

**S4 Table. Continuous functions of the FOS-GRN**.

## Funding

ERAB and JCMG acknowledge the support from UNAM-DGAPA-PAPIIT IN211721 “Patrones genéricos y sistémicos de la diferenciación y la proliferación en los nichos de células troncales: Raíz de Arabidopsis thaliana como sistema de estudio teórico-experimental.” and CONACYT FORDECYT-PRONACES 194186/2020 “Biolog’
smatemática y computacional de sistemas médicos: modulación preventiva de la emergencia y progresión de enfermedades crónico-degenerativas”, respectively.

## Acknowledgments

This work is presented in partial fulfillment towards José Eduardo Chairez-Velóz doctoral degree in the program “Doctorado en Control Automático, del Departamento de Control Automático del Centro de Investigación y de Estudios Avanzados del Instituto Politécnico Nacional”.

## Author contributions statement

Conceived and designed the experiments: ERAB JCMG JECV. Performed the experiments: JECV. Analyzed the data: ERAB JCMG JECV. Contributed reagents/materials/analysis tools: JDV ECCH. Wrote the paper: ERAB JECV JCMG.

## Notes

### Competing Interest Statement

The authors have declared no competing interest.

